# Automated population identification and sorting algorithms for high-dimensional single-cell data

**DOI:** 10.1101/046664

**Authors:** Benedict Anchang, Sylvia K. Plevritis

## Abstract

Cell sorting or gating homogenous subpopulations from single-cell data enables cell-type specific characterization, such as cell-type genomic profiling as well as the study of tumor progression. This highlight summarizes recently developed automated gating algorithms that are optimized for both population identification and sorting homogeneous single cells in heterogeneous single-cell data. Data-driven gating strategies identify and/or sort homogeneous subpopulations from a heterogeneous population without relying on expert knowledge thereby removing human bias and variability. We further describe an optimized cell sorting strategy called CCAST based on Clustering, Classification and Sorting Trees which identifies the relevant gating markers, gating hierarchy and partitions that define underlying cell subpopulations. CCAST identifies more homogeneous subpopulations in several applications compared to prior sorting strategies and reveals simultaneous intracellular signaling across different lineage subtypes under different experimental conditions.

## I. INTRODUCTION TO THE TYPE OF PROBLEM IN CANCER

High-dimensional single-cell data pose challenges in terms of identifying and sorting of cells for investigating tumor heterogeneity. Sorting cells for downstream analysis relies not only on identifying the clusters but also on the gating strategy, which is defined by the gating markers, thresholds and sequence. For manual gating at the FACS machine, typical gating strategies are organized like a family tree. For example, from mature bone marrow cells, lymphocytes are gated from the parent cells and from that gate, T-cells or B-cells are gated, and from those gates, specific T-cell and B-cell types are gated [1]. In particular, sorting out T-cells is equivalent to isolating a CD4+/CD8+ population; the user would first isolate the lymphocytes, then derive the CD3+ cells and from there, would draw a gate around the CD4 positive and CD8 positive subpopulations. This approach assumes prior knowledge of the underlying set of markers that define cell types, the gating hierarchy and relative boundaries for isolating pure cell subpopulations of interest. Selecting these parameters based solely on literature and human perspective introduces bias and variability and could result in contamination among the cell subpopulations.

## II. ILLUSTRATIVE RESULTS OF APPLICATION OF METHODS

CCAST is among a number of newer generation gating algorithms that have been developed for single-cell analysis as the conventional approach of manual, sequential exploration using bivariate dot plots of single cells is not scalable and fails to capture high-dimensional relationships. CCAST is a data-driven sorting strategy that combines any clustering algorithm with silhouette measures to identify underlying homogeneous subpopulations, and then applies recursive-partitioning techniques to generate a decision tree that defines the gating strategy. CCAST produces an optimal strategy for cell sorting by automating the selection of gating markers, the corresponding gating thresholds and gating sequence; all of these parameters are typically manually defined. In the study by Anchang *et al*. [2], CCAST was applied to sort pure subpopulations in both normal bone marrow and breast cancer single-cell data. A closely related automated gating algorithm by Aghaeepour *et al*. [3] called RchyOptimyx uses dynamic programming and optimization techniques from graph theory to construct a cellular hierarchy, providing a qualitative gating strategy to identify target populations to a desired level of purity. This approach provides multiple gating strategies for gating targeted populations, whereas CCAST aims to find the optimal gating strategy [2,3]. Fig. 1 shows the output from CCAST on triple negative SUM159 breast cancer cells from the study by Anchang *et al*. [2] showing the gating steps for 9 subpopulations. Other automated population identification algorithms require some form of manual investigation of their output for population characterization. These include Spanning-tree Progression Analysis of Density-normalized Events (SPADE), a density-based algorithm for visualizing single cell data and inferring cellular hierarchies among subpopulations of similar cells. SPADE has been applied to numerous single-cell flow and mass cytometry datasets addressing a variety of important biological questions from identifying new hematopoietic cell types [1,4] to extensive analysis of cell type specific intracellular signaling following a variety of stimuli including small molecule inhibitors [5]. t-distribution stochastic neighborhood embedding (t-SNE) based algorithms such as viSNE [6] for single-cell visualization and ACCENSE [7] for an Automatic Classification of Cellular Expression by Non-linear Stochastic Embedding) do not require the specification of the target number of unknown clusters. viSNE in particular has been used for population identification in single-cell leukemia data [6]. With the exception of CCAST and RchyOptimyx, these other methods, in addition to previous approaches that include SamSpectral [8], Flowclust [9] and FLAME [10] do not automate the gating or sorting process. More recently, the FlowCAP-II project [11] compared the accuracy and reproducibility across several gating algorithms in terms of identifying cell clusters. All the above methods have some form of clustering algorithm for identification of subpopulations, as a major component. The usefulness of applying CCAST lies in its unbiased optimization of gating and sorting strategies.

**Figure 1.**
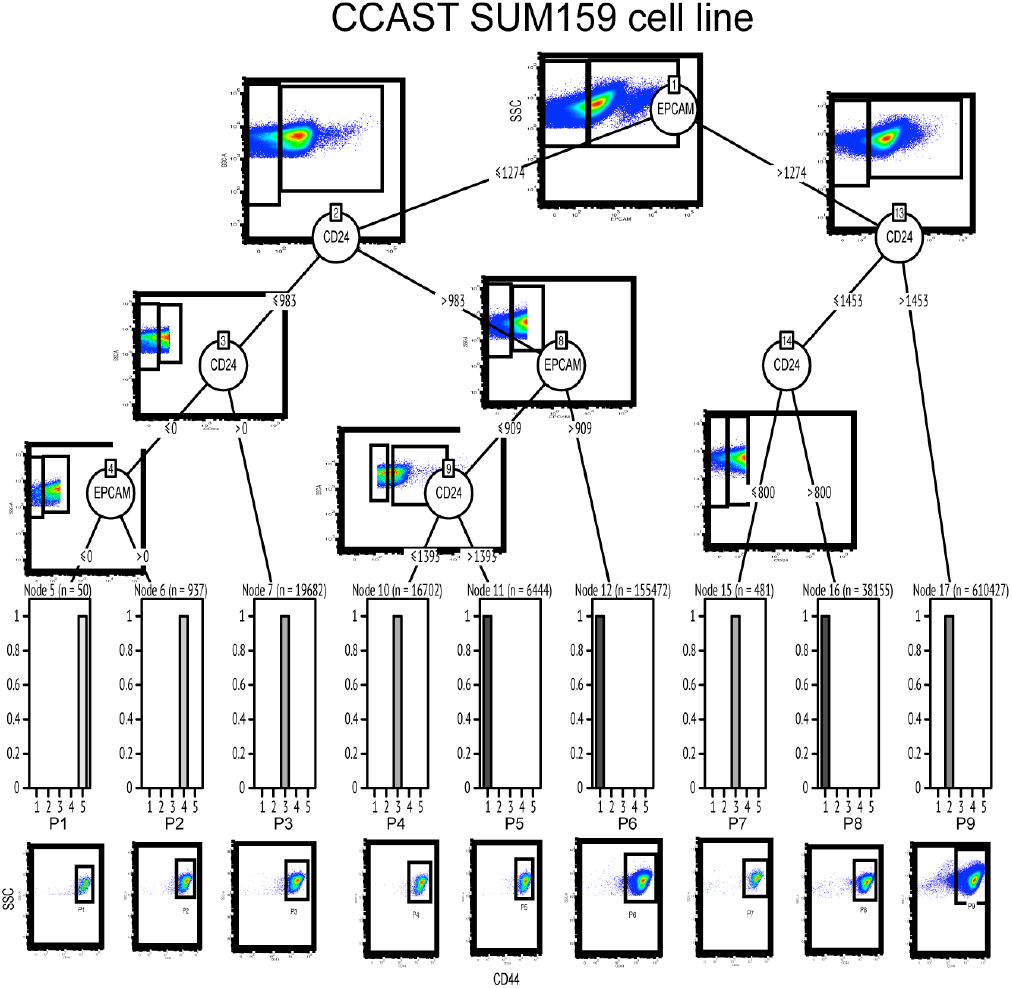
CCAST gating strategy for SUM159 breast cancer cell line

## III. QUICK GUIDE TO CCAST

CCAST formalizes the gating process of single cells as a statistical model and provides a simple unbiased hierarchical 2D gating scheme with the relevant set of marker cut-offs for gating all homogenous cell subpopulations from given Flow Cytometry (FCM) data. Details of the algorithm is summarized in the flowchart of Fig. 2 alongside an analysis of simulated data. Briefly stated, starting with a population of single cell data (Fig. 2A), CCAST performs a cell-clustering algorithm to identify groups of similar cells (Fig. 2B). The clustering can be performed in a variety of ways. CCAST uses a nonparametric mixture model [12] denoted as “npEM” or hierarchical clustering (HCLUST) [13]. Once the clusters are established, CCAST derives a gating strategy that is represented by a decision tree [14] (Fig. 2C), where the nodes specify the gating markers and their thresholds on the edges. The leaves of the decision tree represent the final gated subpopulations. Often the final number of gated populations is greater than the number of initial cell clusters. When this happens some of the populations capture cells from only one cluster, but others capture cells from multiple clusters. For the subpopulations which contain multiple clusters, all or some of the cells in those subpopulations can be removed and CCAST can be retrained on the remaining population, producing a more robust gating strategy because it is less influenced by “contaminating” cells (Fig. 2D). The final decision tree can be used for cell sorting (Fig. 2E) or data analysis (Fig. 2F).

**Figure 2.**
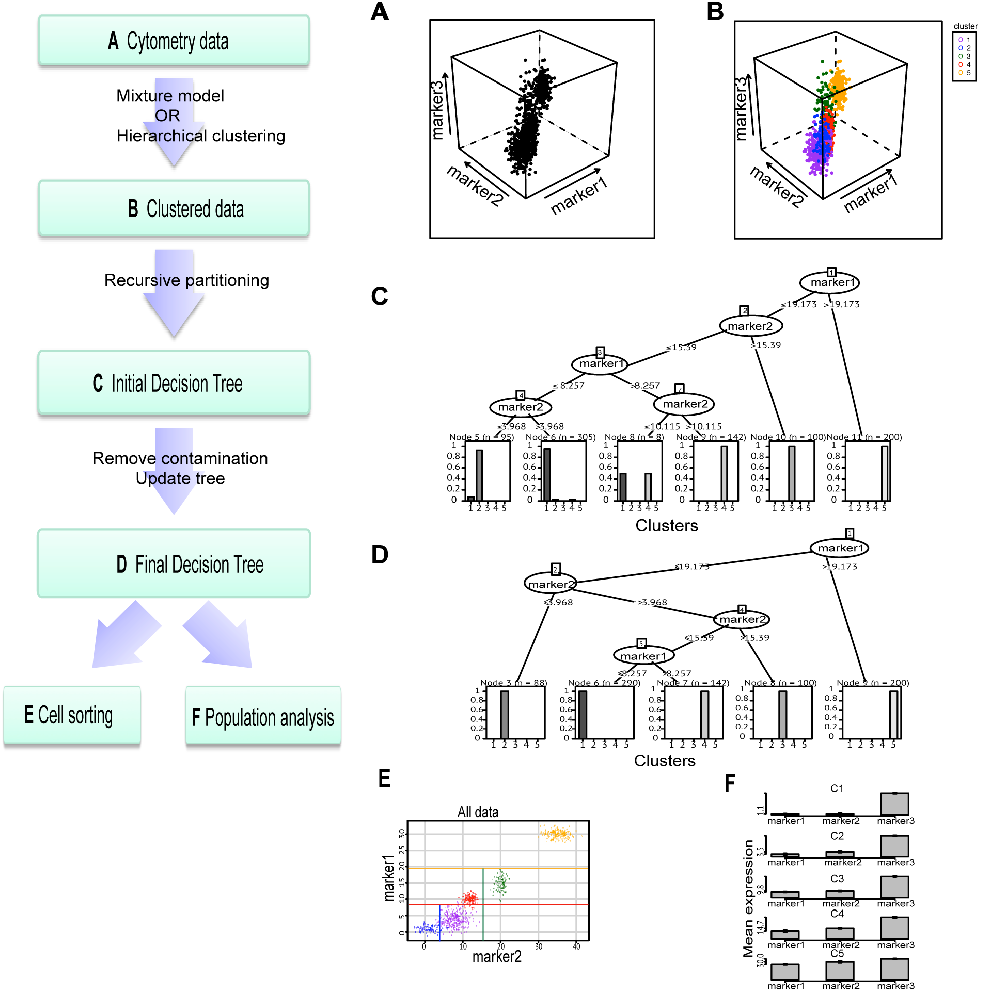
CCAST framework and analysis of simulated data

### A. Other useful settings for CCAST

Although CCAST is optimized for cell sorting and produces a reproducible sorting strategy, it can be used to automatically match homogeneous subpopulations across different experimental conditions for downstream analysis.

